## Supplementary information for "Formation of synthetic RNA protein granules using engineered phage-coat-protein -RNA complexes"

#### Supplementary Results

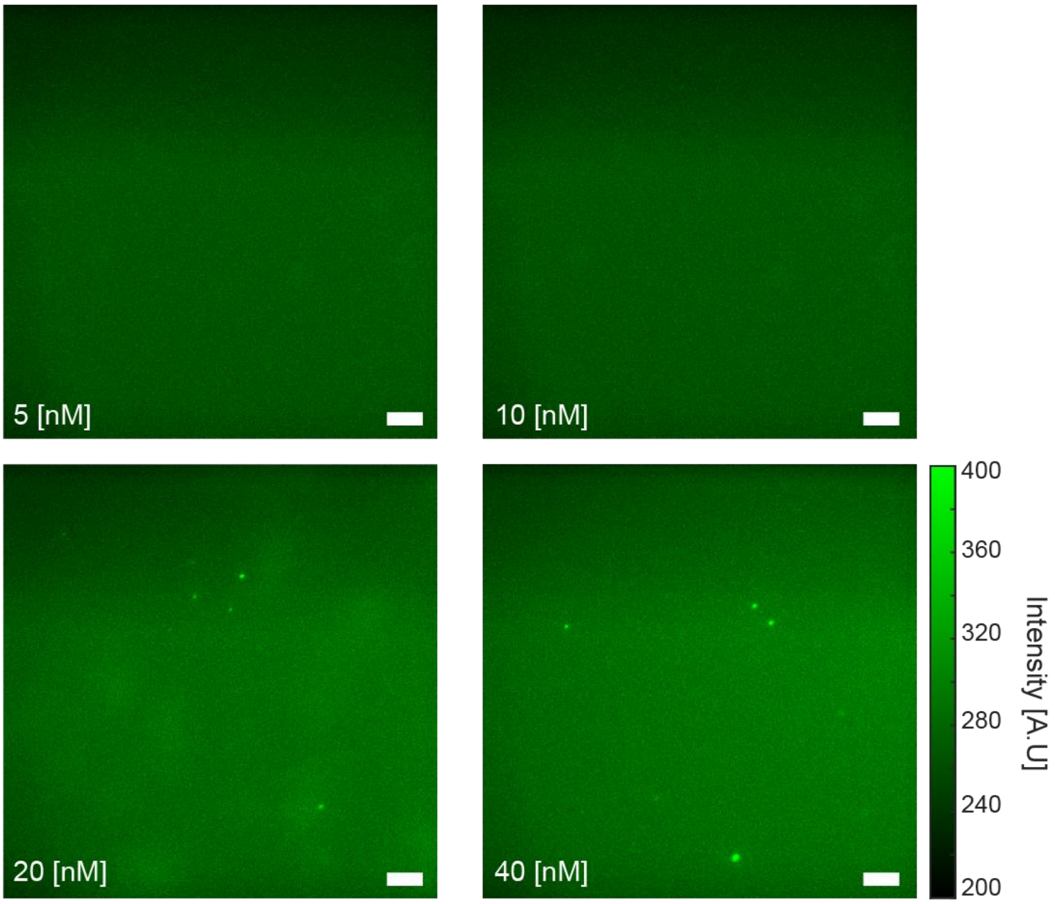


**Supplementary Figure 1**: **PCP-3x titration experiments**. Sample microscopy images of granules reactions with different concentrations of PCP-3x slncRNA. The images show the formation of weak granules at concentration of 20 nM, and brighter, larger granules at 40 nM. Concentration written on the images. Scalebars are 10 µm.

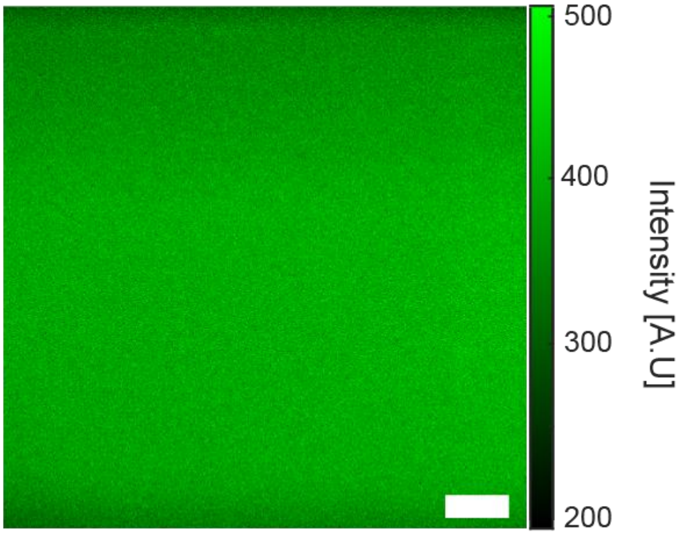


**Supplementary Figure 2**: **Negative control slncRNA only granule reaction**. Scalebar is 10 µm.

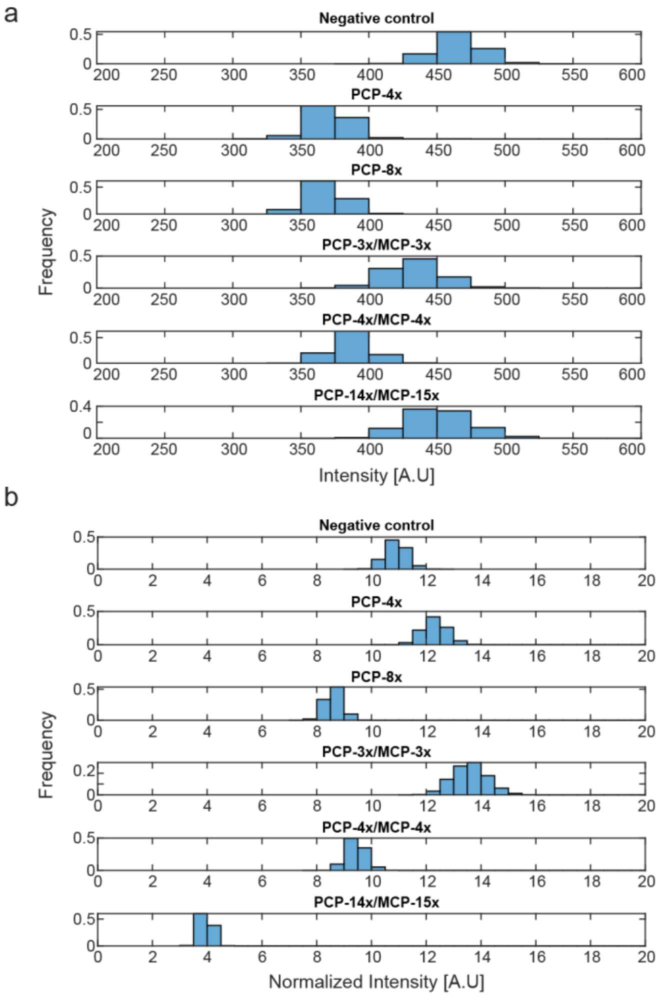

**Supplementary Figure 3**: **Comparison between the background levels from different slncRNA reactions. a,** Absolute values as measured from the microscope. **b,** Values normalized by the estimated number of labeled uracil bases on each slncRNA molecule. Source data are provided as a Source data file.

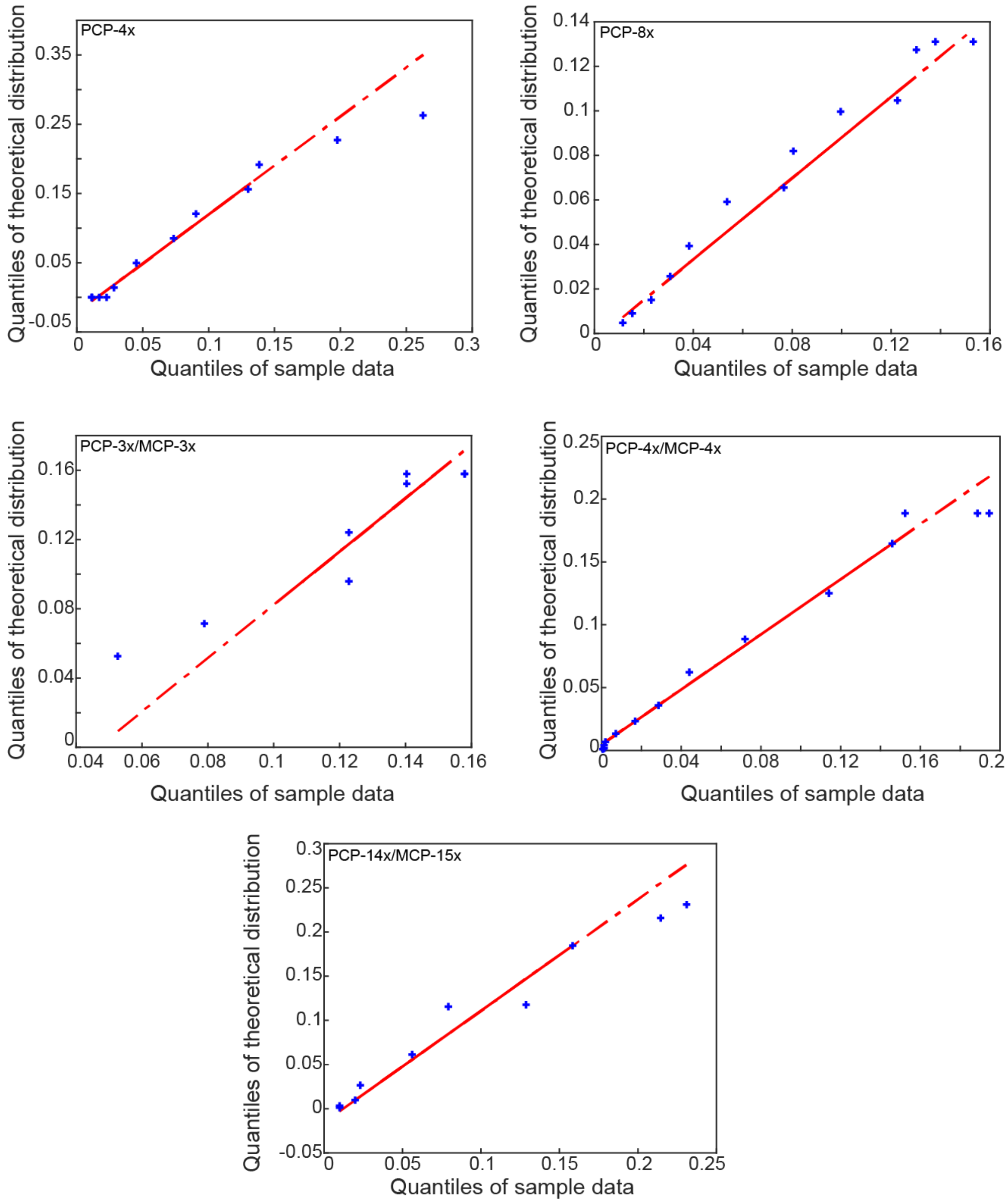


**Supplementary Figure 4**: **QQ-plots of modified Poisson fits**. Quantile-quantile (QQ) plots showing agreement between sample data (experimental observations) and the theoretical Poisson distribution for the fits shown in figure 1f.

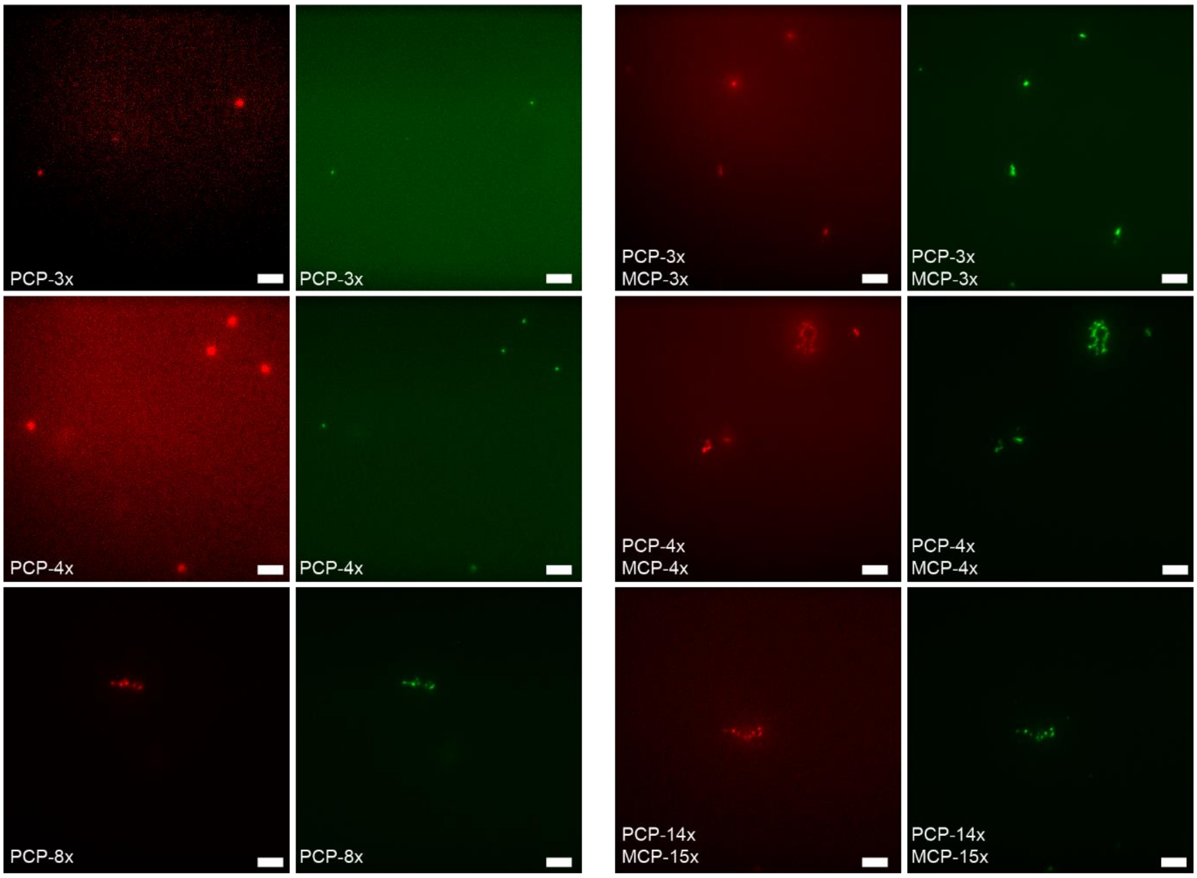


**Supplementary Figure 5**: **SlncRNA-protein granules microscopy images**. Images of slncRNA-protein granules showing the 585 nm channel (red) and the 488 nm channel (green) separately. Overlaid images appear in Figure 2b. All scale bars are 10 µm.

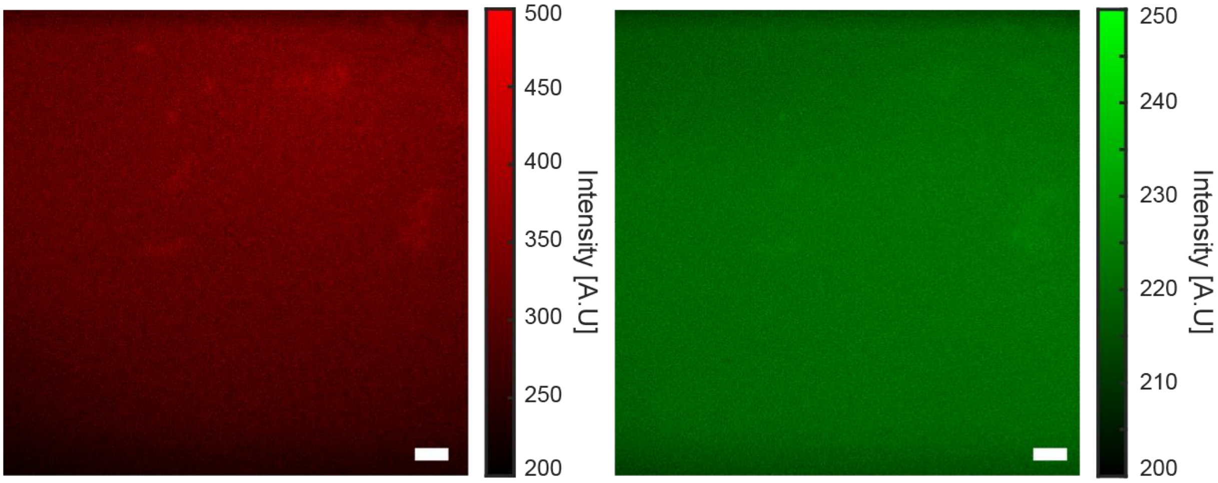


**Supplementary Figure 6**: **Negative control slncRNA-Protein granule reaction.** Scalebars are 10 µm.

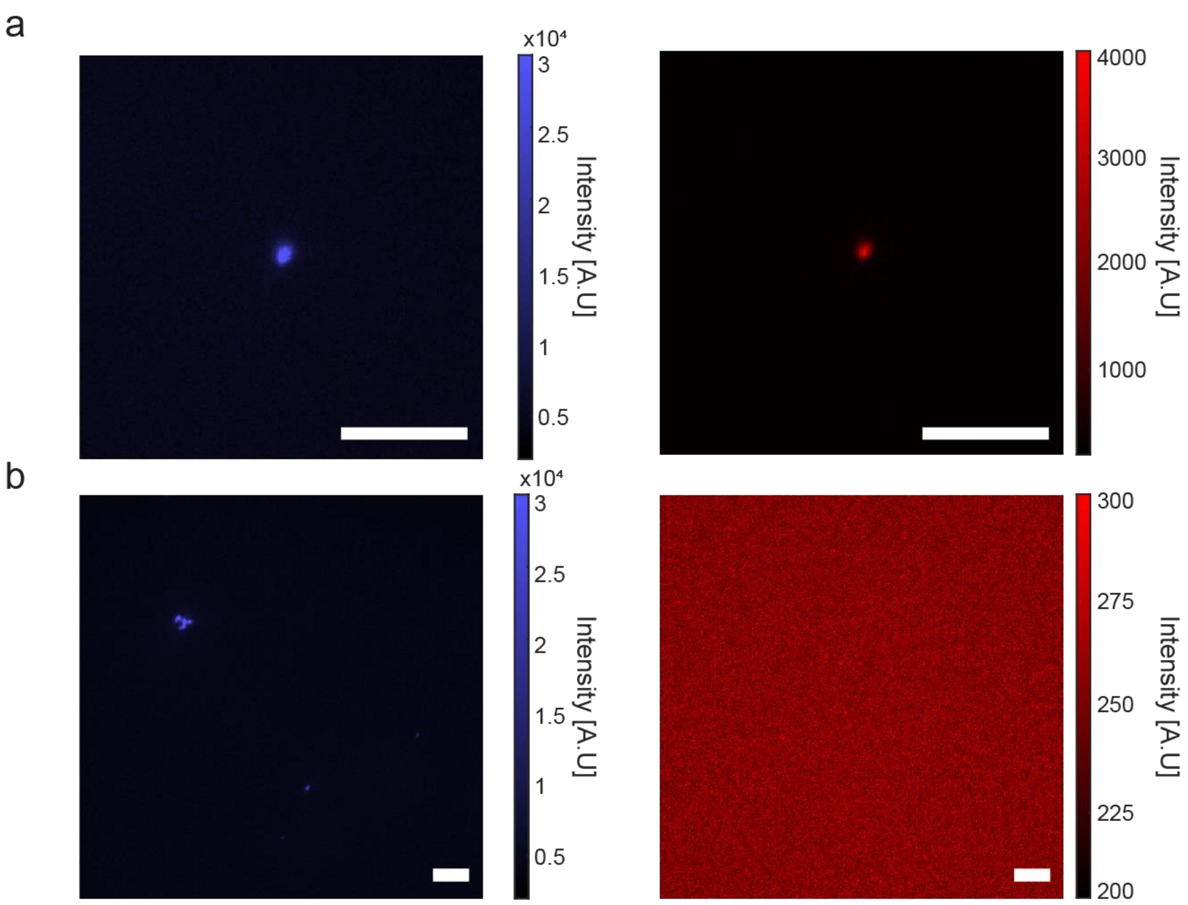


**Supplementary Figure 7**: **Protein affinity experiment.** a, Granule formation reaction with class II slncRNA (PCP-3x/MCP-3x) labeled in AF405 labeling (left), and tdMCP-mCherry (right), showing colocalization between the protein and the RNA. b, Granule formation reaction with class I slncRNA (PCP-8xlabeled in AF405 labeling (left), and tdMS2-mCherry (right), showing RNA structures with no corresponding protein fluorescence. Scalebars are 10 µm.

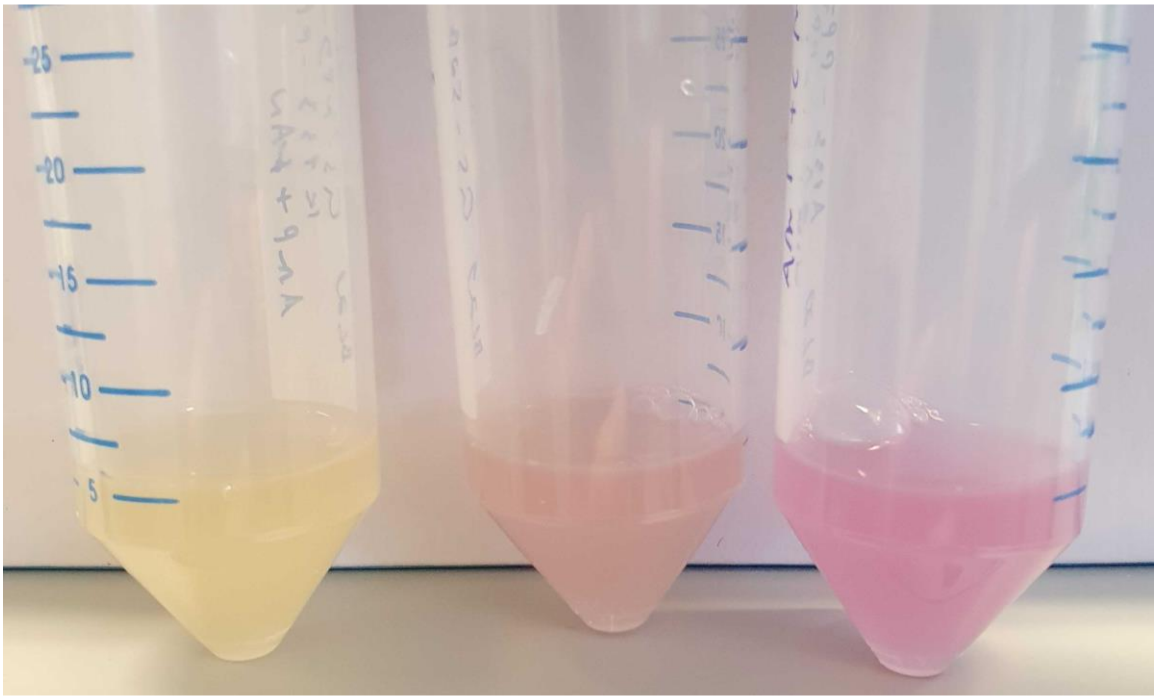


**Supplementary Figure 8**: **Bacterial cultures images.** From left to right: *E. coli* BL21-DE3 expressing tdPP7-mCherry with the negative control slncRNA. *E. coli* BL21-DE3 expressing tdPP7-mCherry with PCP-4x/QCP-5x slncRNA. *E. coli* BL21-DE3 expressing tdPP7-mCherry with PCP-24x slncRNA.

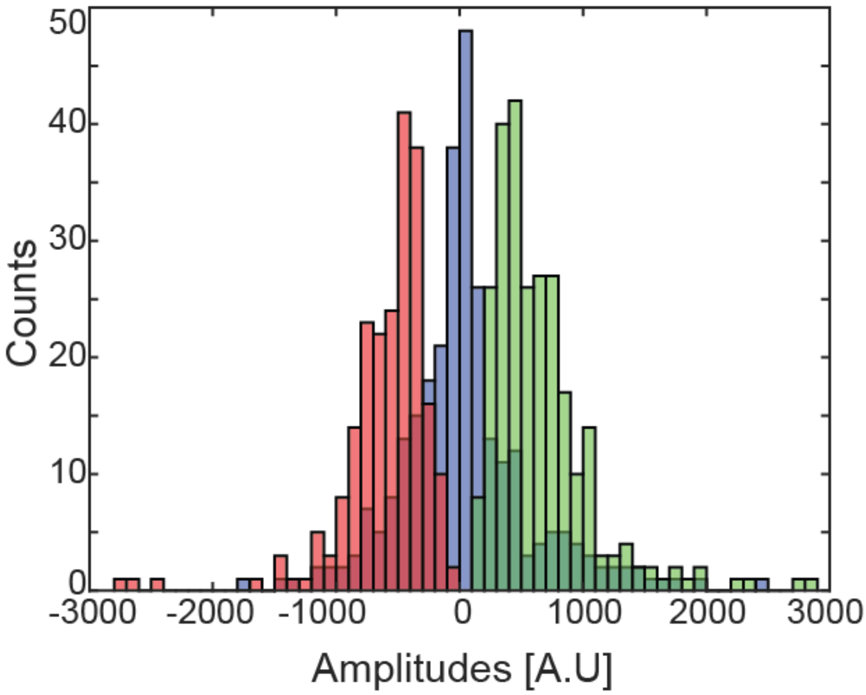

**Supplementary Figure 9**: **PCP-24x granules amplitude distribution**. **a**, Empirical amplitude distributions gathered from 391 traces *in vivo* from cells expressing the PCP-24x slncRNA together with the tdPCP-mCherry protein. Green – Positive amplitudes (insertion events), red – negative amplitudes (shedding events), blue – unclassified events. Source data are provided as a Source data file.

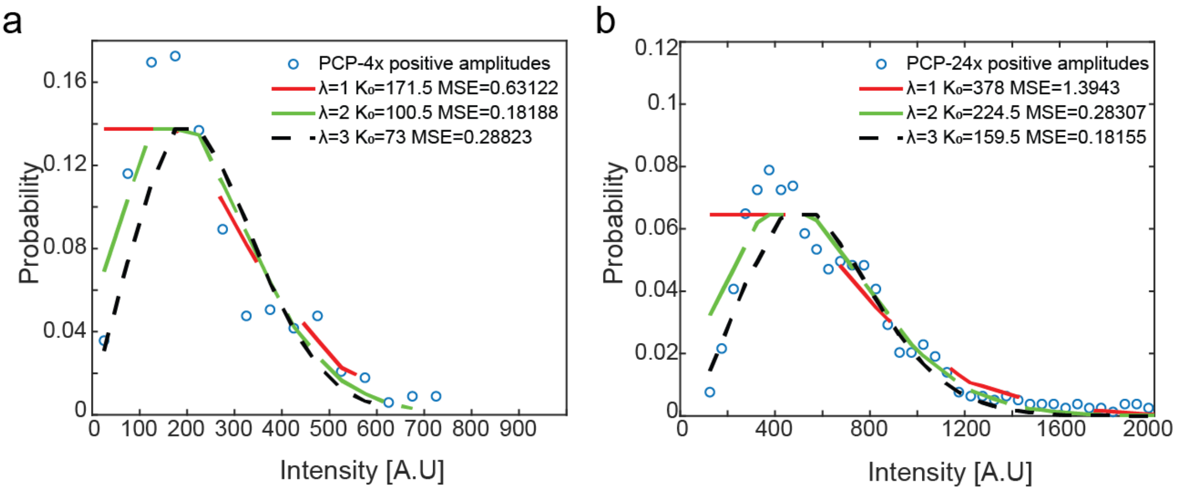


**Supplementary Figure 10**: **Fitting of amplitude data to Poisson distributions.** (a-b) Poisson functions fits for the amplitude distribution of insertion events assuming 1, 2, or 3 mean events (λ values). MSE values represent mean squared error between the empirical distribution and the theoretical modified Poisson functions. **a**, Data collected from 255 PCP-4x/QCP-5x signal traces. **b**, Data collected from 391 PCP-24x signal traces.

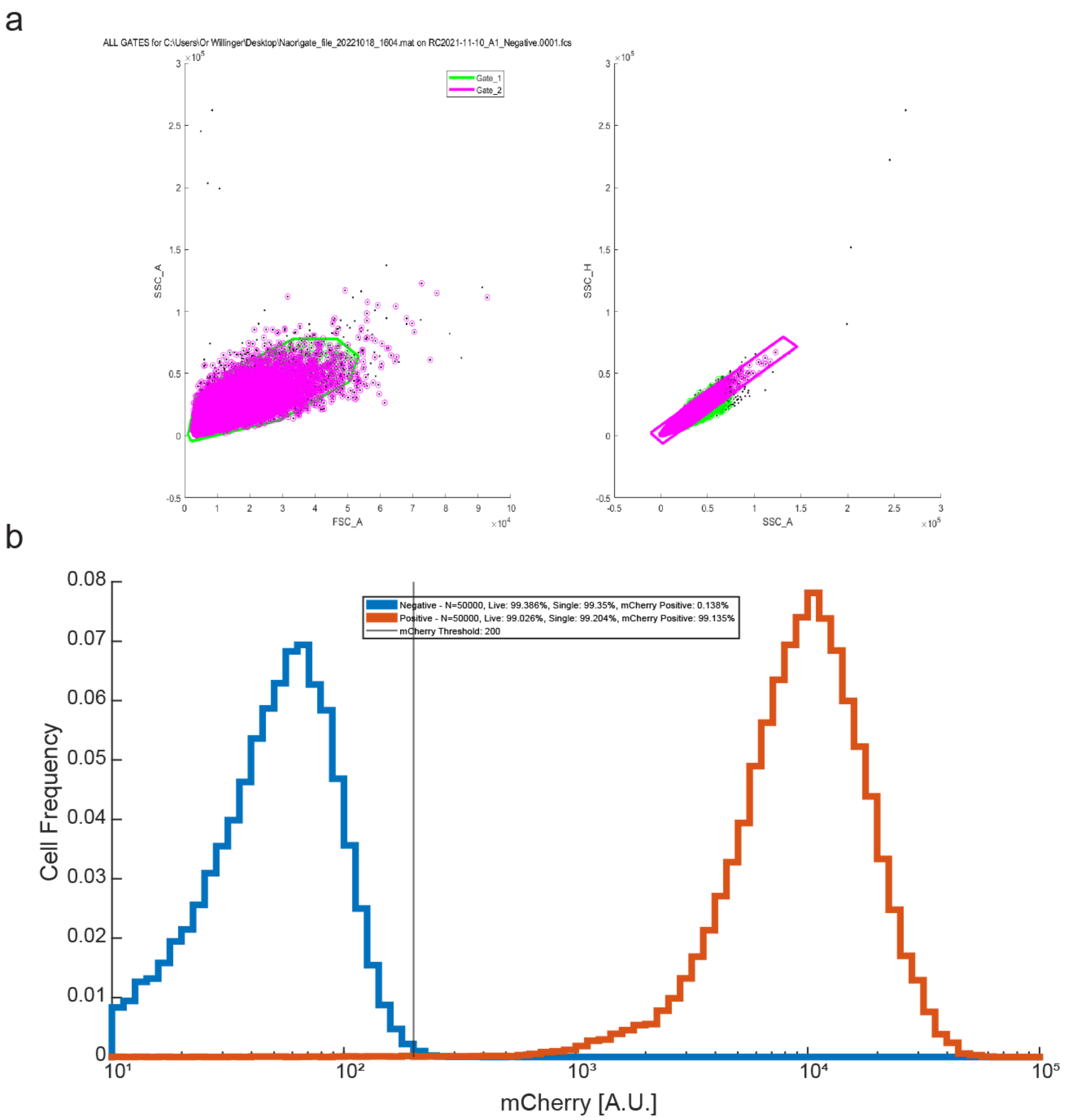


**Supplementary Figure 11**: **Sample FACS gating strategy. a,** Example graphs representing the gating strategy on the SSC-A and FSC-A parameters during cell sorting for a sorting experiment. (left) Live cells gating, (right) single cell gating. **b,** A plot of the mCherry data for a single sorting experiment showing a positive population (expressing tdPCP-mCherry) in orange, and a negative population in blue.

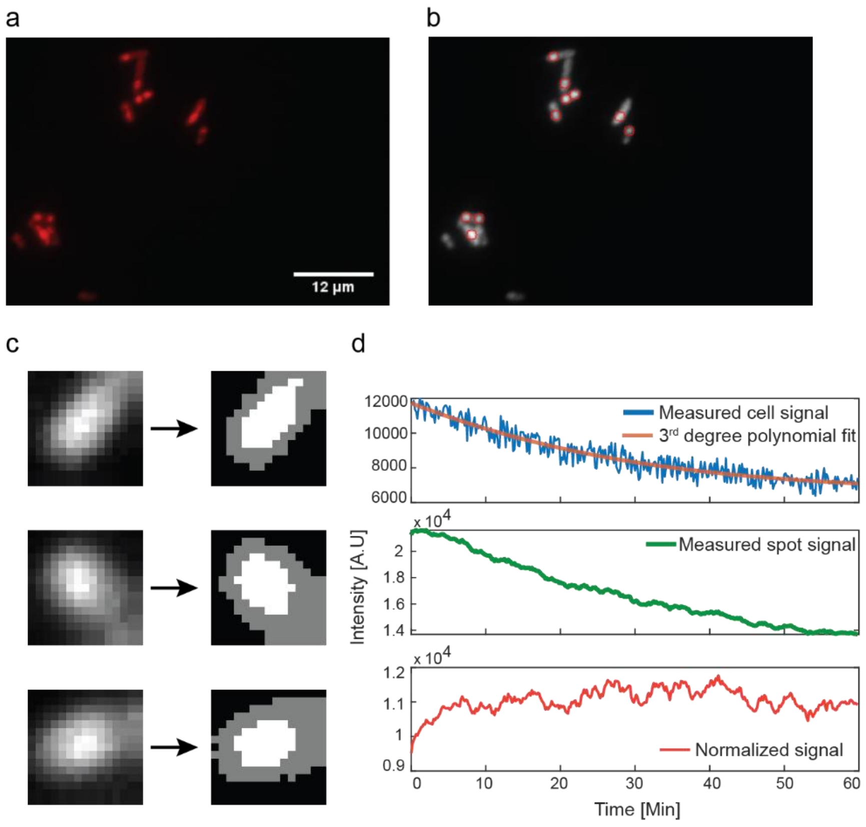


**Supplementary Figure 12**: **Image processing scheme**. **a**, Raw microscopy image showing bacterial cells containing bright spots. **b**, Bright spots are identified and their position over time and space is recorded. **c**, The environment of each spot is classified into 3 regions, based on intensity values. The brightest pixels are classified as 'spot' (marked in white), the darkest pixels are classified as 'dark background' indicating empty space, and pixels with intermediate values are classified as cell background (marked in gray). **d**, The mean values of the spot pixels and cell background pixels are recorded over time resulting in the spot signal (green) and cell signal (blue). The spot signal is then normalized to remove photobleaching and global background effects (red).

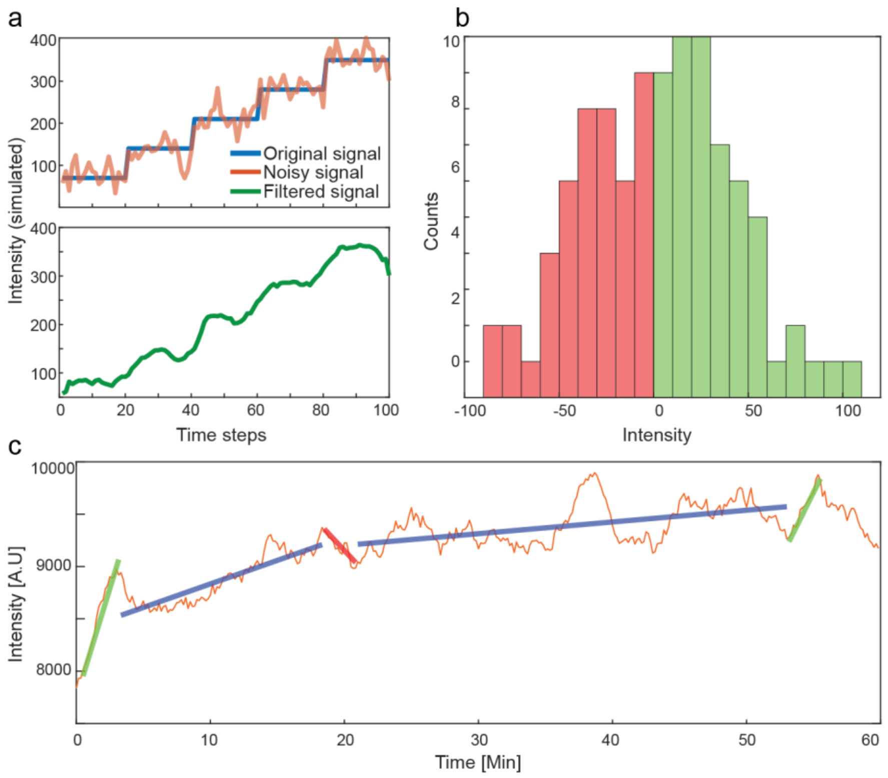


**Supplementary Figure 13**: **Identification of burst events**. **a**, Top: simulated step signal (blue) with added white Gaussian noise (orange). Bottom: noisy signal after moving average filter. **b**, Intensity difference distribution for the signal presented in panel A. **c**, Sample experimental signal (orange) overlaid with markers indicating identified segments in green, blue, and red, corresponding to positive bursts, quiescent segments, and negative bursts.

### Supplementary text

#### Numerical simulations of signal types

To check that our analysis is consistent with an underlying random burst signal, we simulated three types of base signals with added noise components. For each simulation type, 1000 signals of 360 time-points were simulated and analyzed using the same data analysis process described in the methods section.

We simulated flat constant signals, gradually ascending signals, and signals containing multiple burst events. Two noise components were added to all signals, based on our noise model. White Gaussian noise of magnitude 40 [A.U] peak-to-peak amplitude, matching the value estimated from experimental traces, and an exponential component, simulating photobleaching (Supplementary Fig. 14).

We then applied our burst-detection algorithm described above and found that for the flat signal (Supplementary Fig. 14 a) positive and negative bursts (green and red respectively) and non-classified events are detected. However, a closer examination of the results reveals that the burst amplitude width is smaller by a factor of ~5-10 as compared with the experimental data bursts, and the total number of events observed (458 positive, 452 negative, and 298 non-classified segments found) is significantly smaller than the experimental data, indicating roughly 1 event per signal, as expected from our base assumption that a rare noise event occurs once in a thousand time points. For the gradually increasing signal with additional noise, (Supplementary Fig. 14 c) a negligible number of negative burst-like events was detected by our algorithm, with a pronounced bias towards positive events (1111 positive, 9 negative and 467 non-classified). The scarcity of events can be explained by the positive bias in the signal which results in a steep increase in the statistical threshold for event identification. Similar simulations with a decreasing signal show a mirror image of amplitude distribution (data not shown).

Finally, a signal designed to mimic our interpretation of the experimental data containing randomly distributed instantaneous bursts, both increasing and decreasing with multiple possible amplitudes was analyzed (Supplementary Fig. 14 e). Our simulated signals resulted in a symmetric amplitude distribution, comprising of non-Gaussian or skewed amplitude distributions. Additionally, the range of amplitudes observed is 2-3x larger as compared with the case for the constant signal, with the non-classified amplitudes presenting a wider distribution. A total of 2298 positive, 1831 negative and 2489 non-classified segments were found.

#### Estimating statistical significance of burst events in all traces recorded

To compute whether the number of burst events identified via our algorithm is statistically significant, we simulated a constant base-line intensity amplitude with overlaid white Gaussian noise. For each numerical trace, we simulated 360 times points (corresponding to a ~60-minute experimental trace) and identified the total number of “increasing” and “decreasing” burst events in accordance with the algorithm described in detailed above. Here, we used m = 10 (see eqn. 1.8) consecutive increasing or decreasing instantaneous signal difference events as our threshold. We identified 458 and 298 increasing and decreasing burst events respectively in 1000 simulated traces with constant baseline. By comparison, we found 2298 and 1831 increasing and decreasing burst events respectively in 1000 simulated traces containing bursts, which using Fisher's test yield a p-value of 4e-309 and 2e-310 for the significance of the increasing and decreasing burst findings.

We repeated this statistical test for experimental data, comparing the PP7-4x data against traces measured from cell containing only tdPP7-mCherry with no expression of our RNA cassettes, using the latter as a baseline akin to the constant signal simulations. We identified 7 increasing and 6 decreasing burst events in 150 traces gathered from the cells lacking RNA binding sites, while for the PP7-4x data we identified 112 increasing and decreasing burst events in 255 experimental traces, which using Fisher's test yields a p-value of 2e-13.

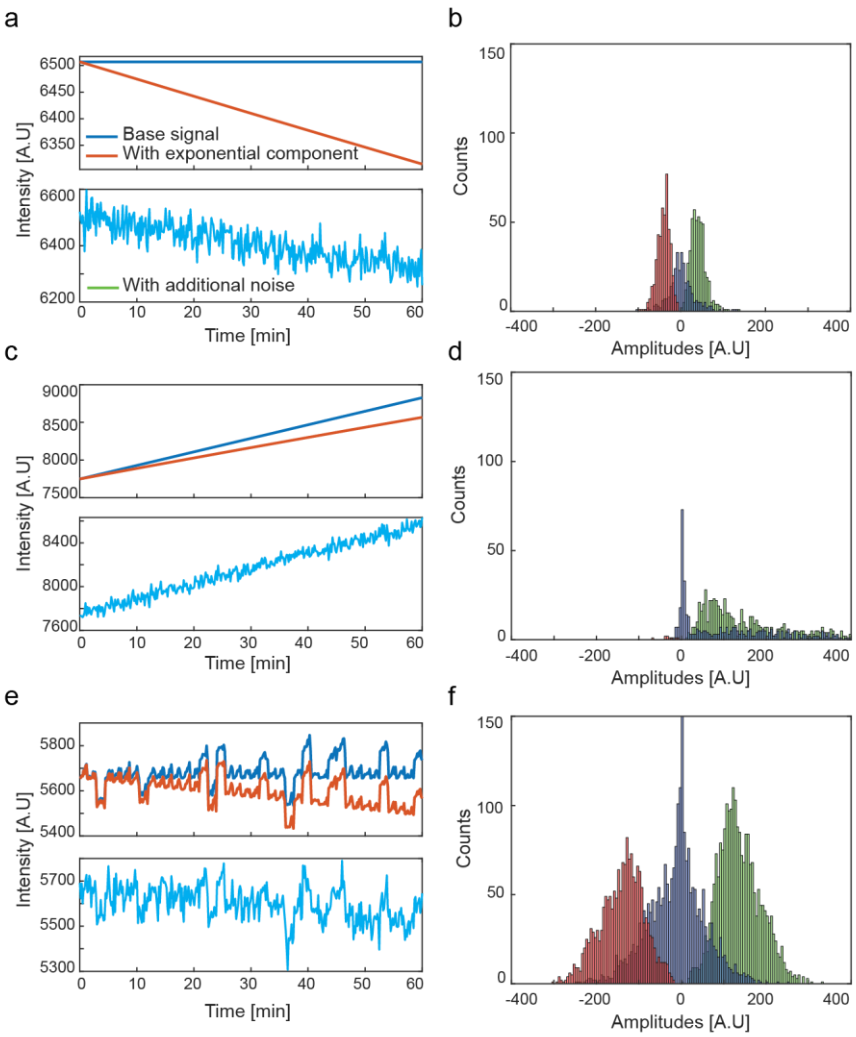


**Supplementary Figure 14**: **Signal type simulations**. **a**, Simulated constant signal (blue), with photobleaching (orange), and added noise (cyan). **b**, Amplitude distributions of burst events identified from 1000 constant signals. **c**, Simulated signal with slope (blue), with photobleaching (orange), and added noise (cyan). **d**, Amplitude distributions of burst events identified from 1000 sloped signals. **e**, Simulated signal with burst events (blue), with photobleaching (orange), and added noise (cyan). **f**, Amplitude distributions of burst events identified from 1000 bursty signals.

#### Signal analysis parameter selection

##### Subframe length

As part of the analysis process of the *in-vivo* microscopy experiments, the immediate surroundings of each discovered bright spot are recorded as a sub-frame containing the spot at its center, from this sub-frame the mean spot intensity and mean background intensity are calculated. The selection of the sub-frame length used to calculate the background intensity is an important parameter in the analysis process that might bring about unwanted noise into the resulting statistics when analyzing *in vivo* images. A large sub-frame might include other cells, with possibly different bright spots of themselves, inserting a bias into both the cell background intensity, and spot intensity signals. On the other hand, a small sub-frame might not have a sufficient spot-to-background area ratio, resulting in an underestimated cell background signal.

To select the appropriate sub-frame length, we analyzed the PCP-24x data with sub-frames of different lengths – 10, 14, 20, and 30 pixels. Supplementary Fig. 15 a shows an example of this where the orange, yellow and red squares correspond to sub-frames of 10,14 and 20 pixels in length, and the panel itself constitutes a 30-pixel wide sub-frame. The criteria for this selection process are the mean ratio between cell area to spot area; percentage of frames where this ratio is less than one; and the ratio between the spot mean intensity to the cell mean intensity without any filtering or fitting. These criteria are designed to find the length that does not cause an overestimation of cell background against spot or vice versa (as could be the case where more than one bright spot fall inside the sub-frame). From these tests we learned that lengths of 10 and 14 pixels result in a mean ratio of less than two (i.e., on average the sizes of the bright spot and of its surrounding environment are equal) (Supplementary Fig. 15 b). However, a sub-frame length of 10 pixels results in nearly a fifth of frames where the cell background is less than one and thus potentially underestimated (Supplementary Fig. 15 c). Finally, the intensity ratios show that the mean ratio does not vary much between the different options, however the spread is more conserved for lengths of 10 and 14 pixels (Supplementary Fig. 15 d). Following these tests, we chose a sub-frame length of 14 pixels for our analysis process.

##### Moving average span

The moving average window span is a critical component in the signal analysis process. It is used both as a noise reduction filter, and as a means to bias sharp signal jumps. The filter span plays another significant role, as it is the minimal allowed length for a burst duration. Choosing a small value might introduce false positives into the statistics, while a large value would cause many actual burst events to be discarded. To find the optimal span length we compare the number of events found in a simulated flat signal, such a signal should not produce any bursts under noise-less conditions. For this we simulated 1000 constant signals, 360 time points each, with an added white Gaussian noise and an exponential component and applied our data analysis procedure. (Supplementary Fig. 16 a). An ideal result for this test would be less than one event of each type, i.e., positive, and negative bursts, per signal (Supplementary Fig. 16 b).

We further show that using intermediate span length values (9-13 time points), has little effect on the qualitative nature of the results (Supplementary Fig. 16 c, d).

Following these tests, we decided on a span of 13 time points. This value results in one event or less of each type per simulated signal, while still allowing us to record the statistical nature of the experimental signals.

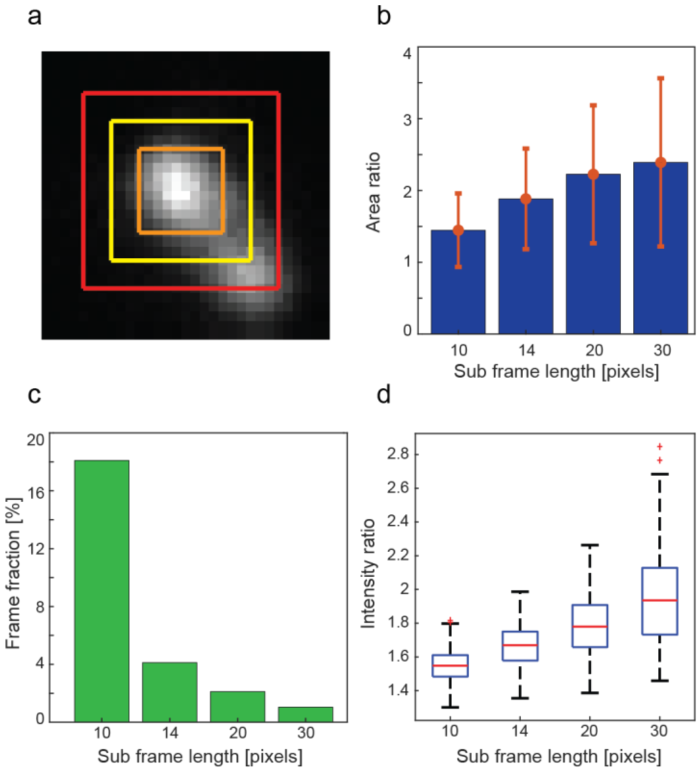


**Supplementary Figure 15: Subframe length selection**. **a**, Example of different sub-frame lengths. Image is a sub-frame with length of 30 pixels. Red, yellow, and orange squares correspond to sub-frames of length 20, 14 and 10 pixels accordingly. **b**, Mean ratio between cell background area to spot area (both are in number of pixels), calculated from 200 sub-frames of bright puncta inside bacterial cells. **c**, Percentage of cells where the area ratio presented in (b) is less than one, indicating probable underestimation of the cell background. **d**, Ratio between spot mean intensities to cell background mean intensities (i.e., each spot is divided by its corresponding cell background) calculated from 200 bright puncta inside bacterial cells. On each box, the central mark indicates the median, and the bottom and top edges of the box indicate the 25th and 75th percentiles, respectively. The value for 'Whisker' corresponds to ±1.5 IQR (interquartile rate) and extends to the adjacent value, which is the most extreme data value that is not an outlier. The outliers are plotted individually as plus signs. Source data are provided as a Source data file.

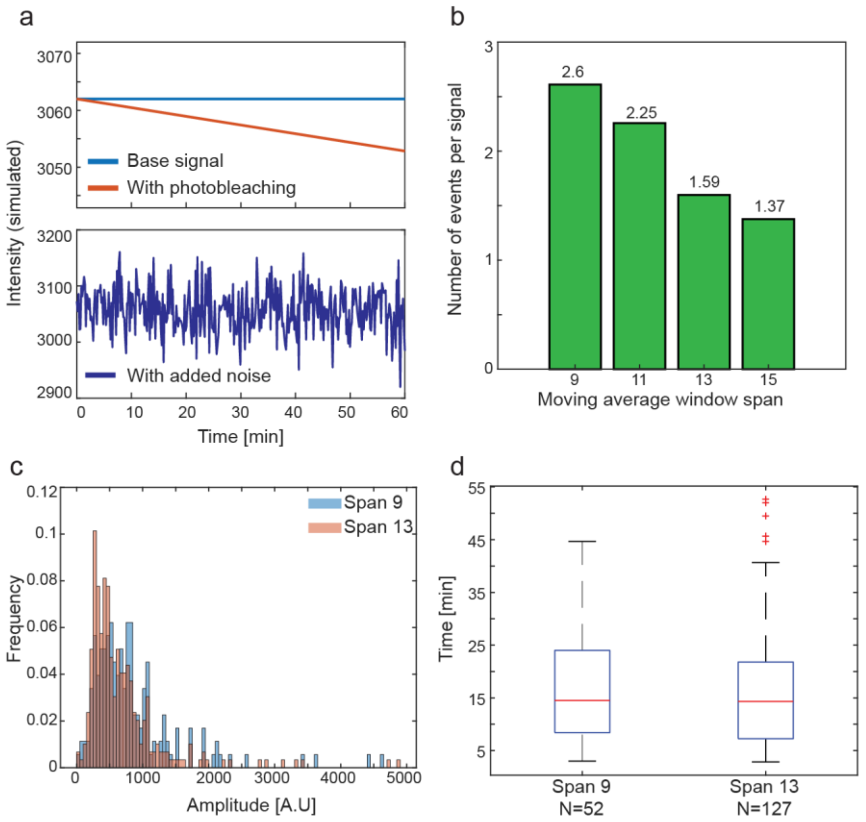


**Supplementary Figure 16: Moving average span length selection**. **a**, Sample simulated signal used for testing. Blue line is the underlying constant signal, orange line represents the same signal with an added photobleaching component. Cyan signal is the orange signal with added white Gaussian noise. **b**, Total number of identified events of any kind per simulated signal. **c**, Positive amplitude histograms of PCP-24x data analyzed using a moving average filter of 9 time points (blue) and 13 time points (orange). **d**, Duration between positive events of PCP-24x data analyzed using a moving average filter of 9 time points and 13 time points. On each box, the central mark indicates the median, and the bottom and top edges of the box indicate the 25th and 75th percentiles, respectively. The value for 'Whisker' corresponds to ±1.5 IQR (interquartile rate) and extends to the adjacent value, which is the most extreme data value that is not an outlier. The outliers are plotted individually as plus signs. Source data are provided as a Source data file.
